# Identification of Viral Variants from Functional Genomics Data

**DOI:** 10.1101/2025.01.31.635891

**Authors:** Florian Röckl, Caroline C. Friedel

## Abstract

The analysis of virus knockout mutants is a common approach for studying the role of individual viral genes in viral infections and is increasingly performed using functional genomics sequencing experiments, e.g. RNA-seq or ATAC-seq, of infected cells. Identifying viral variants directly in these experiments avoids additional genome sequencing and allows confirming the presence of particular mutations directly in the experiment of interest. Here, we present a pipeline to directly identify viral variants from these functional genomics data. This combines existing SNP callers with novel methods for identifying deletions, insertions, and corresponding inserted sequences. The latter address the problem that existing structural variant callers performed poorly on functional genomics data with large variations in coverage. We evaluated the pipeline on RNA-seq data for infection with wild-type Herpes simplex virus 1 (HSV-1) and null mutants of important HSV-1 proteins. Comparison of variants identified by our pipeline with the descriptions of the original publications showed that we could correctly recover the introduced mutations. Thus, our pipeline offers researchers a fast and easy way to verify the existence of variants in the viral genome without additional genome sequencing experiments.

**Availability:** The pipeline is implemented as a workflow for the workflow management system Watchdog and is available at https://github.com/watchdog-wms/watchdog-wms-workflows/ (Variant-CallerPipeline).

## Introduction

Advances in molecular biology and genetics provide new technologies for studying virus infections and the role of individual viral genes during infection. This provides the basis for the development of treatments against virus infections or for their use as tools in genetic engineering, vaccine development, or gene therapy. A common approach is the creation of mutant virus strains (see e.g. (Johnston and McFadden, 2004)) containing single nucleotide polymorphisms (SNPs) or deletions or insertions (indels) of sequences that disturb functions of individual viral genes. For well-studied viruses like herpesviruses, such experiments have been conducted for decades. Accordingly, many mutant strains have been generated decades ago, often before complete genome sequences of these viruses were available (e.g. in Marsden et al. (1976); Post and Roizman (1981); Davison et al. (1984); Stow and Stow (1986); Perry et al. (1986); DeLuca and Schaffer (1987, 1988); Fenwick and Everett (1990); Smith et al. (1992)). These were commonly passed between laboratories and used for a multitude of experiments. The precise genome location of mutations or inserted sequences are often poorly documented and other non-documented mutations may have been introduced either with the original mutation or in the time since. Furthermore, even for more recently created mutants, it is important to verify presence of mutations, in particular, if results from experiments do not meet expectations. The standard approach to identify mutations in viral genomes is genome sequencing, which requires separate (potentially repeated) genome sequencing experiments. Due to advances in high-throughput sequencing technologies, however, analyses of virus gene functions are now commonly performed using sequencing-based functional genomics assays of virus-infected cells, such as RNA-seq, ATAC-seq or ChIP-seq (e.g. in Djakovic et al. (2023)). Since these generally also provide nucleotide coverage of viral genomes, they provide the unique opportunity to identify viral variants directly in the experiment of interest without additional genome sequencing.

In this article, we present a pipeline to automatically identify viral variants in functional genomics data of virus infections, including SNPs, deletions and insertions and (optionally) inserted sequences. This pipeline uses existing SNP calling methods, in particular bcftools (Li, 2011) and VarScan (Koboldt et al., 2012)), which we found to perform well also for RNA-seq or other functional genomics data that exhibit large variations in read coverage across the viral genome (see e.g. Figure S1). In contrast, existing structural variant callers (in particular DELLY (Rausch et al., 2012), GRIDSS2 (Cameron et al., 2021), and BreakDancer (Chen et al., 2009)) performed poorly in identifying insertions and deletions in viral null mutants from these data. This is likely because they expect approximately uniform read coverage as in genome sequencing. We thus implemented a new approach to identify deletions and insertions based on gaps in read coverage and clipped (i.e. partial) read alignments. We combined this with *de novo* assembly using rnaSPAdes (Bushmanova et al., 2019) to identify inserted sequences. Analysis of RNA-seq data for wild-type and null mutant herpes simplex 1 (HSV-1) infection shows that our pipeline allows fast and easy identification of viral variants and their precise genomic locations to better characterize null mutants generated decades ago.

## Methods

The virus variant caller pipeline was implemented as a workflow for the workflow management system Watchdog and is available at https://github.com/watchdog-wms/watchdog-wms-workflows/ (Variant-CallerPipeline). The deployment of required software is performed with conda (https://conda.io, using conda-forge and bioconda channels) using the deployment functionality of Watchdog. Watchdog also supports easy parallelization of workflow runs on computing clusters and monitoring of workflow execution, which can be made use of when running our pipeline. The workflow takes as input read alignments against the viral genome in BAM format for one or more virus-infected samples and read sequences in FASTQ format (only necessary for detection of inserted sequences). We used BWA (Li and Durbin, 2009) for read alignment as it is very fast and requires little memory, but any read alignment program can be used that provides SAM/BAM output, includes read sequences in the output and produces clipped read alignments if only parts of a read can be aligned to the viral genome. Notably, since we are not interested in identifying splicing junctions, there is no need to use a splicing-aware aligner for RNA-seq data. Example input files can be found at https://doi.org/10.5281/zenodo.14266852.

The variant caller pipeline is divided into two main parts, which are described in the following: (1) SNP calling and (optionally) strain identification and (2) indel detection and (optionally) identification of inserted sequences.

### SNP calling

Figure 1 provides an overview of the steps performed for SNP calling. First, the variant callers bcftools (Li, 2011) and VarScan (Koboldt et al., 2012) are applied independently to each input BAM file. Both tools provide the identified SNPs in the variant call format (VCF) (Danecek et al., 2011). Next, so-called *consistent* SNPs are selected if they are identified by both bcftools and VarScan. If more than one replicate is available, SNPs are considered consistent if they are detected by both tools in all replicates. Consistent SNPs are then mapped to viral features, e.g., genes, coding sequences, or introns, given a gene annotation in GTF format for the viral genome.

**Figure 1.**
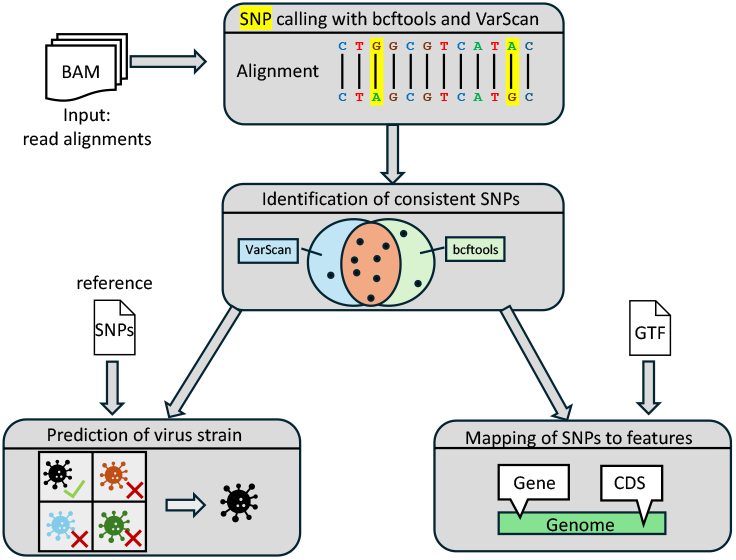
Overview of the steps the pipeline employs for SNP calling.

Furthermore, if a set of reference SNPs for different virus strains is provided by the user, the pipeline performs a prediction of the virus strains. This is useful both for verifying the virus strain used in the experiment and the parental strain from which a particular null mutant was generated. An example file with reference SNPs for HSV-1 strains 17, F and KOS 1.1 is included with the example input files and the Watchdoch module for strain identification (identifyStrain, available at https://github.com/watchdog-wms/watchdog-wms-modules).

For strain identification, the following distance *D* is calculated for each reference strain: *D* = |*S*_1_ *∪S*_2_| *−* |*S*_1_ ∩ *S*_2_|, with *S*_1_ the set of consistent SNPs identified for the particular virus used in the experiment and *S*_2_ the set of reference SNPs for a particular strain. This measure is largely independent of the reference genome sequence used for read alignment as the strains differ predominantly by SNPs and a few small indels. The strain with the smallest distance *D* is then predicted for the virus.

### Indel detection

Insertions and deletions in viral genomes are determined as outlined in Figure 2. First, per-base read coverage, i.e. the number of reads overlapping each genome position, and clipped reads (= reads with unaligned parts) are extracted from each input BAM file using samtools (Li et al., 2009). The results are then used as input for indel calling as outlined below. Identified indels are subsequently also mapped to genomic features. In addition, the pipeline can identify inserted sequences by combining the results from insertion detection with *de novo* read assembly obtained with SPAdes (Prjibelski et al., 2020) if raw read sequences in FASTQ format are provided.

**Figure 2.**
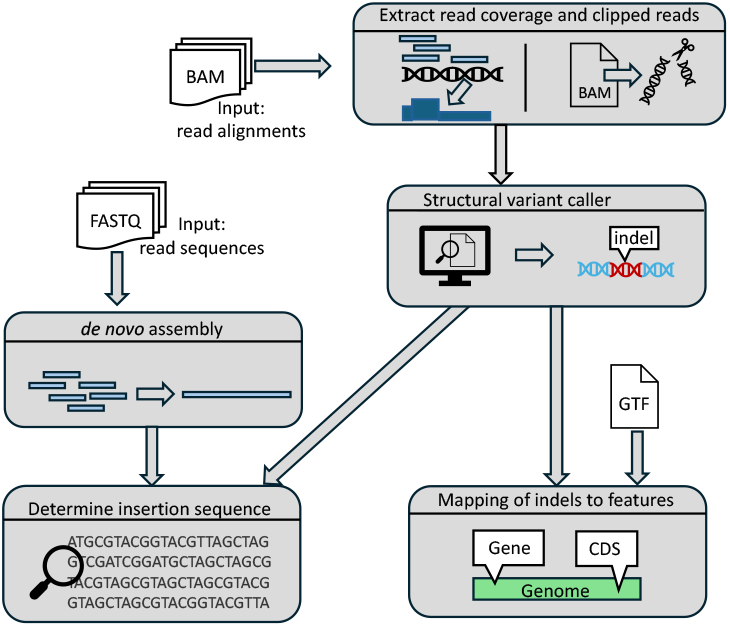
Overview of the steps the pipeline employs for indel identification.

#### Deletion detection

The pipeline first detects potential deletions by identifying regions of the genome with very low read coverage compared to (i) the complete genome using a global threshold and (ii) the surrounding genomic regions using a local threshold. For this purpose, a global z-score is calculated for each position, comparing the logarithm of the read coverage (= log read coverage) for this position to the mean and standard deviation for the complete genome. If this is below a stringent global threshold, the position is labeled as a potential deletion. If it passes only a less stringent global threshold, a local z-score is calculated comparing the log read coverage at this position to the mean and standard deviation of the previous *n* nucleotides (nt) before the current position (by default *n* = 500). If the local z-score is below the local threshold, the position will also be labeled as a potential deletion.

The local z-scores are used as read coverage can vary massively between positions in functional genomics data. This is exemplified for an RNA-seq sample in Figure S1. However, calculating local z-cores for every position, which requires calculating the mean and standard deviation over the preceding *n* nt for every genomic position, is very costly. Thus, the stringent global z-score threshold is first employed to identify clear-cut cases of potential deletions. Local z-scores are only calculated for less clear-cut cases. Optionally, a user-defined length threshold can also be used to exclude very short deletions.

#### Deletion verification

Candidate deletions are subsequently verified using clipped read alignments. As depicted in Figure 3, reads crossing a deletion in the genome can only be aligned with gaps to the reference genome. If the alignment is performed using a non-splice-aware read aligner, such as BWA, this will result in clipped read alignments where only parts of the read are aligned to the genome. This can also occur with splice-aware read aligners as the start and end nucleotides of the deletion commonly do not match canonical splicing signals expected by many splice-aware read aligners.

**Figure 3.**
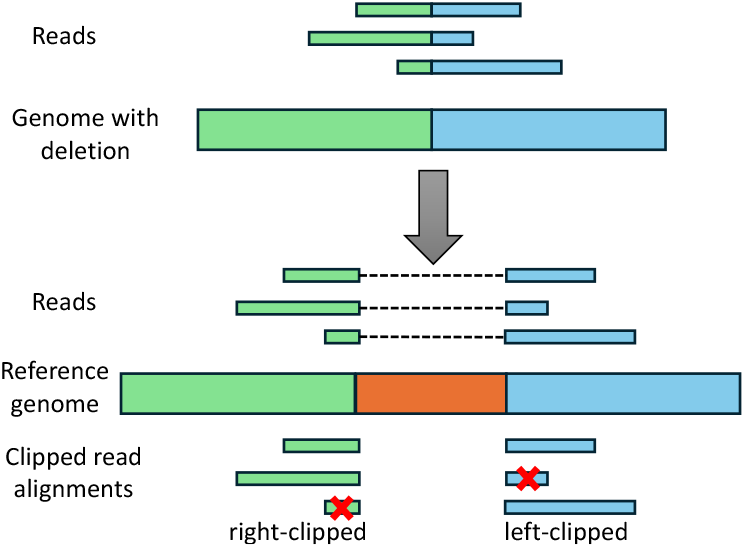
Illustration of clipping at deletion sites. The top shows the mutated viral genome that contains the green and blue sequences from the reference genome below, but the orange sequence was deleted. Reads from the mutant viral genome can thus only be aligned partially to the reference genome, resulting in clipped read alignments (at the bottom). If both parts of the read are sufficiently long to be aligned to the genome, this will result in multiple (clipped) alignments per read. If a part of the read is too short for alignment (marked by a red cross), this part will not be aligned at all.

Thus, deletions should exhibit a peak of right-clipped reads ending at the deletion start and a peak of left-clipped reads beginning at the deletion end. Such peaks of clipped reads are again identified using both a global and local z-score, which are separately calculated for peaks of right-clipped and left-clipped reads. For the global z-score at each position, the number of clipped reads is compared against the mean and standard deviation for the same type of clipped reads across the whole genome. For the local z-score, the number of clipped reads is compared against the mean and standard deviation for a window starting *x* nt upstream of the candidate peak and ending *x* nt downstream of the candidate peak (by default *x* = 20), excluding the peak position itself. If both the global and the local z-scores pass a global (default: 10) and local (default: 50) threshold, respectively, the position is considered a peak. In addition, a minimum number of reads is required for a peak (default: 10 reads).

To verify deletions, the pipeline identifies pairs of right-clipped and subsequent left-clipped peaks (i.e. the *clipping pattern* of deletions) and determines whether the positions of the two peaks overlap with a deletion detected based on the per-base read coverage. Subsequently, the clipped sequences of the corresponding clipped read alignments (i.e. the unaligned part of the read in this alignment) are extracted from the BAM file and position-weight-matrices (PWMs) are computed from the sequence profiles of the clipped sequence parts. As can be seen in Figure 3, the PWMs of the clipped sequence parts on either side of the deletion should match the reference sequence on the opposite side of the deletion.

To test this, the best match of each PWM is determined in a window around the opposite deletion end. The score of a potential match is calculated as the sum of log-odds scores over all positions comparing the value of the PWM for the nucleotide at this position against the background probability of that nucleotide in the complete genome sequence. The best match for a PWM is the match with the highest score. If the best matches for both PWMs have a score >0 or at least one has a score that may contain an insertion. This special case was observed for one of the data sets analyzed in the results section. In this case, the potential insertion sequence is determined as described in the next section and can be further analyzed.

It should be noted that our approach for predicting deletions may also identify splicing events in RNA-seq data if a non-splice-aware aligner is used for read alignments. However, splicing is rare in viruses and the few cases detected can easily be excluded after mapping the deletions to the genome annotation.

#### Insertion detection

Our pipeline also uses clipped read alignments to determine insertions since reads containing part of an inserted sequence cannot be completely aligned to the genome (see Figure 4). Originally, we expected that the resulting clipping pattern should consist of a peak of right-clipped reads at a genome position *n* preceding the insertion followed by a peak of left-clipped reads at position *n* + 1. However, the examples of null mutants created by insertions that we investigated showed a different pattern, consisting of peak of left-clipped peaks at position *n* and a peak of right-peaked reads at position *n* + 1 (see Figure 4). This results from the first and last position of inserted sequences matching the genome on the other side of the insertion and is likely a consequence of the use of homologous recombination for inserting sequences (Bollag et al., 1989).

**Figure 4.**
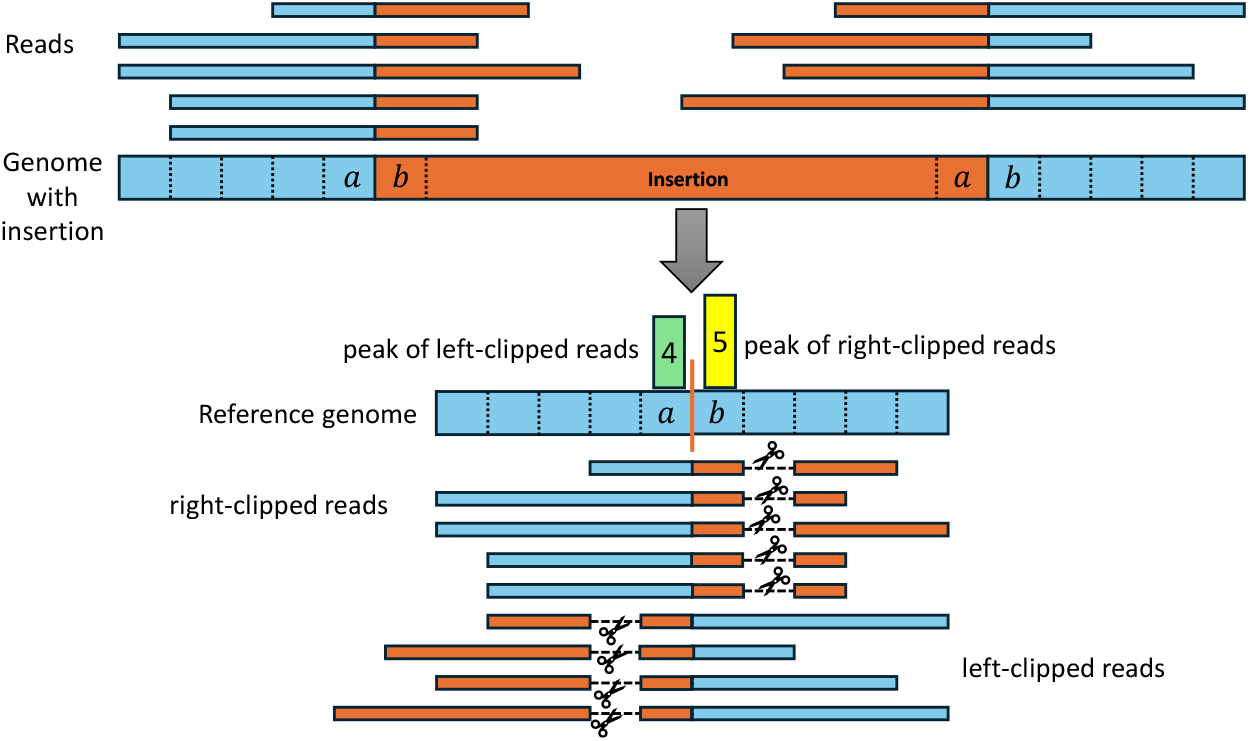
Illustration of read clipping at insertion sites. The top shows the mutated viral genome (blue) that contains an inserted sequence (orange) not present in the reference genome. Reads spanning the boundary of the insertion therefore contain parts of both the reference and insertion sequence. When aligned to the reference genome, the part of the reads containing the insertion sequence (orange) have to be clipped since they cannot be aligned to the reference genome. We observed that commonly the start and/or end of the insertion also matches the reference genome directly before and after the insertion site (in this example, 1 nt matches on each side). As a result, reads can be aligned beyond the insertion site, resulting in a distinctive insertion clipping pattern with a peak of left-clipped positions one or more positions left of a peak of right-clipped positions.

To allow for such matches between the insertion start and/or end to the surrounding genomic regions, we introduced a parameter *ϕ* determining the maximum number of such matches that are allowed. Thus, any pair of positions for a left-clipped peak *p*_*l*_ and a right-clipped peak *p*_*r*_ is used to predict an insertion if *p*_*r*_ *− p*_*l*_ + 1 *≤ ϕ*. In the example in Figure 4, *p*_*r*_ *− p*_*l*_ + 1 = 2. For each identified insertion, we extract the non-aligned parts of clipped reads to calculate consensus sequences for the insertion start and end, respectively. These consensus sequences are commonly 30-40 nt long.

To identify the remaining central part of the inserted sequences, the pipeline optionally performs a *de novo* sequence assembly using rnaSPAdes (Bushmanova et al., 2019), a modification of the genome assembler SPAdes (Prjibelski et al., 2020) for application to RNA-seq data. Assembly is performed for all reads, which may also include non-viral sequences. Following this, the consensus sequences for the insertion start and end are aligned to the resulting assembled contigs using BWA. If a match for both consensus sequences is found, the assembled sequence starting with the consensus of the insertion start and ending with the consensus of the insertion end is extracted. Insertion sequences containing only one of the consensus sequences are also extracted but are flagged for special attention. The origin of the inserted sequences can then be confirmed using BLAST (Altschul et al., 1990).

It should be noted that we also investigated using only *de novo* assembly for detection of both insertions and deletions by aligning the assembled contigs to the reference genome (see results). However, this either resulted in too few or too many indels depending on parameters, thus we did not pursue this approach for the pipeline.

### Results

### Input data

We applied our pipeline to previously published 4sU-seq data for infection with wild-type HSV-1 strain 17 and null mutants of multiple HSV-1 proteins (Wang et al., 2020). 4sU-seq is a variant of RNA-seq based on sequencing newly transcribed RNA obtained by 4-thiouridine (4sU) RNA tagging (Windhager et al., 2012). The 4sU-seq data included null mutants for the following HSV-1 proteins:

- ICP4, created by a SNP, which resulted in a temperature sensitive mutant (TsK) (Marsden et al., 1976; Davison et al., 1984)
- ICP0 and ICP22, created by deletions (Stow and Stow, 1986; Perry et al., 1986; Post and Roizman, 1981)
- ICP4, ICP27 and *vhs*, created by insertions (DeLuca and Schaffer, 1988, DeLuca and Schaffer, 1987; Smith et al., 1992; Fenwick and Everett, 1990).

Two replicates were available for all viruses, except for the ICP4 knockout by insertion (ΔICP4), for which only one replicate was available.

In addition, we analyzed RNA-seq data for human brain organoids (Rybak-Wolf et al., 2023) infected with an HSV-1 strain 17 virus engineered to express green fluorescent protein (GFP)(Snijder et al., 2012). Here, RNA-seq data was available for brain organoids from two genetically distinct induced pluripotent stem cell lines, each infected for 3 and 6 days (2 replicates each, resulting in 8 samples).

All 4sU-seq and RNA-seq data were aligned against the HSV-1 strain 17 genome (GenBank accession: JN555585) using BWA and then fed into the pipeline. The performance of the pipeline was evaluated by comparing the results with the descriptions of the original publications. The insertion sequences that were extracted by the pipeline from the sequence assembly were investigated with the NCBI BLAST webserver to identify their origin.

### SNPs in the TsK mutant

Our pipeline identified 28 consistent SNPs in the TsK mutant, three of which were in the ICP4 gene. One of these was consistent with the sequence change identified by Davison et al. (1984) as responsible for the mutant phenotype: a replacement of a C:G base pair by a T:A base pair that resulted in a substitution of the 475^th^ codon from an alanine by a valine codon. Our pipeline matched this missense mutation to a SNP at nucleotide 129,708 and thus for the first time described the precise genomic location of this mutation. It furthermore showed that the TsK mutant different from its parental strain 17 by an additional 27 other SNPs, whose effect remains unclear. In particular, one of the other two SNPs identified in ICP4 leads to a second amino acid change in ICP4 from serine to asparagine.

### Deletions identified for HSV-1 null mutants

To detect deletions in the 4sU-seq data, our pipeline was run with a stringent global z-score cut-off of -2.5, a less stringent global cut-off of 0.0 and a local z-score cut-off of -6.0. No minimum length was required for the deletions. This resulted in the identification of the deletions shown in Table 1. Here, a deletion found in ΔICP0 and a deletion found in ΔICP22 infection matched the target gene and approximate length described in the corresponding articles (Stow and Stow, 1986; Perry et al., 1986; Post and Roizman, 1981). Furthermore, the sequences found directly up- and down-stream of the predicted deletions matched the target sequences of the restriction enzymes used in the corresponding experiments to create the deletions (XhoI & SalI for ΔICP0; PvuII & BstEII/Eco91I for ΔdICP22). Thus, we could recover the exact locations of the introduced deletions.

**Table 1.** Deletions detected by the pipeline on the 4sU-seq data and whether this represents the deletion described in the original papers describing the null mutant, a deletion in the parental strain or a known intron. For the known intron in ICP22, the position of the intron is also indicated in the last column. The genes US10-US12 overlap at the deletion position.

| Mutant/strain | Type | Start position | End position | Gene |
| --- | --- | --- | --- | --- |
| ΔICP0 | described deletion | 120913 | 123031 | ICP0 |
| ΔICP22 | deletion in parental strain F | 132276 | 132280 | ICP22 |
| ΔICP22 | described deletion | 133243 | 134072 | ICP22 |
| ΔICP27 | deletion in parental strain KOS 1.1 | 144838 | 144849 | US10;US11;US12 |
| ΔICP27 | known intron | 132404 | 132497 | ICP22 (132375-132543) |
| Δvhs | known intron | 132390 | 132513 | ICP22 (132375-132543) |
| wild-type strain 17 | known intron | 132392 | 132505 | ICP22 (132375-132543) |

In addition, a deletion was identified for wild-type strain 17, ΔICP27 and Δ*vhs* that fell into a known intron in the ICP22 gene. Although this is spliced in all samples, it was not detected in ΔICP4 and TsK infection. This was likely due to the fact that read coverage was much lower for ΔICP4 and TsK infection on the whole genome as ICP4 is necessary for optimal expression of other HSV-1 genes (Watson and Clements, 1980). A further deletion in the 5’ UTR of ICP22 identified in ΔICP22 infection corresponded to a genome deletion in the parental strain F from which the ΔICP22 strain was derived. Similarly, a deletion identified in ΔICP27 infection is already present in its parental strain KOS 1.1.

### Insertions identified for HSV-1 null mutants

Insertions were predicted for the 4sU-seq data using default values. Here, 40 nt around each peak position were used to calculate the local z-score. In addition, we required at least 10 clipped reads for each peak and allowed a maximum overlap *ϕ* of 10 nt for the insertion ends and the surrounding genome regions. The consensus sequences obtained from the clipped parts of the read had to be at least 10 nt long. The identified insertions are listed in Table 2.

**Table 2.** Insertions detected by the pipeline for any of the 4sU-seq data and whether this represents the insertion described in the original papers describing the null mutant or an insertion in the parental strain. Here, the confirmed insertion sequence is Overlap = overlap between the ends of the inserted sequence and the surrounding genome sequences.

| Mutant | Type | Position | Overlap | Gene |
| --- | --- | --- | --- | --- |
| ΔICP4 | described insertion (stop codons, HpaI recognition site) | 130376 | 4 | ICP4 |
| ΔICP27 | described insertion (E. coli lacZ gene) | 113648 | 3 | ICP27 |
| ΔICP27 | insertion in parental strain KOS 1.1 | 140458 | 3 | US5;US6;US7 |
| Δvhs | described insertion (cloning vector with lacZ gene) | 91923 | 2 | vhs |

Here, all but one insertion represented the insertions used to create the null mutant according to the publications (DeLuca and Schaffer, 1988, 1987; Smith et al., 1992; Fenwick and Everett, 1990). The additional insertion identified in ΔICP27 represented a known insertion in the parental strain KOS 1.1. In particular, we could confirm the insertion of lacZ genes in both ΔICP27 and Δ*vhs* infection by BLASTing the predicted insertion sequences obtained from the assembly. For the ΔICP4 mutant, the insertion of a small 16 nt sequence could be directly confirmed from the consensus sequences of the insertion start and end, which overlapped. This insertion sequence contained the 3 stop codons, one for each frame, and a recognition site of the HpaI restriction enzyme described in the original publication.

In all three cases, we are the first to describe the precise genomic locations of these insertions. Interestingly, we found that the position for the insertion in the *vhs* coding sequence (251st codon = in the Δ*vhs* mutant described in the corresponding publication (Fenwick and Everett, 1990) was likely calculated based on a wrong strand assignment. The *vhs* gene is located on the negative strand, with the coding sequence ranging from positions 91167 (stop codon) to 92636 (start codon). Accordingly, the identified insertion position is in the 238th codon. However, if the codon position is erroneously calculated from the positive strand, the insertion would be after the first position of the 252nd codon (excluding the stop codon), which is consistent with the original publication. Fenwick and Everett (1990) also described the insertion site to be in the unique recognition site of the NruI restriction enzyme in the *vhs* gene. The center of this NruI recognition site is at our identified insertion position, thus confirming that it is the correct position. This further highlights the usefulness of our pipeline.

### Insertions in the GFP-expressing HSV-1 virus

According to the original publication describing this virus (Snijder et al., 2012), an EGFP gene with a mouse cytomegalovirus promoter was was inserted between the open reading frames (ORFs) UL55 and UL56 and a LoxP site (= a 34 nt DNA sequence recognized by the Cre recombinase enzyme) was inserted downstream of the UL23 ORF. Two insertion sites at positions 46665 and 116147 were identified in 8 and 7 samples, respectively, located downstream of the UL23 coding sequence and between UL55 and UL56, respectively. The insertion sequence for the first insertion indeed contained a LoxP site while BLAST analysis identified the insertion sequence of the second insertion site to match several cloning vectors containing the GFP gene. Thus, we correctly identified the precise genome positions of both insertions.

It should be noted that the insertion at position 116147 was actually identified as a deletion between positions 116147 and 116154 into which an insertion was placed. This special case is predicted by the pipeline if the PWMs obtained from the clipped reads cannot be matched to the opposite end of the deletion and an insertion sequence can be identified from the assembly. Unfortunately, the description in the original publication on how the sequence was inserted is not sufficiently detailed to explain how this small deletion was generated during the insertion process, but it is most likely a consequence of the experimental approach used.

Additional insertions were identified at positions 62143, 106984, and 119496 in 4-8 of the samples. However, no insertion sequences could be extracted from the assembly for insertions at positions 62143 and 106984 based on the consensus sequences from the clipped reads, while the insertion sequence for 119496 matched the genome downstream of the predicted insertion site. From this and inspection of the genome at this positions, we concluded that these represented artifacts from repetitive sequences. This highlights how the combination of consensus sequences from the clipped read parts and the assembly can be used to filter out incorrectly identified insertions.

### Comparison to *de novo* assembly

For comparison, we also investigated whether deletions and insertions can be identified directly from the contigs assembled by rnaSPAdes instead of performing the analysis of read coverage and clipped read alignments performed by our pipeline. For this purpose, contigs assembled for the 4sU-seq data of HSV-1 mutant infections were aligned against the reference genome using minimap2 (Li, 2018). However, this showed that assembled contigs often contained very small indels (∼1-50bp) compared to the reference genome, which would result in a large number of predicted indels if we included all of them. We, thus included a filtering step to exclude short indels. Furthermore, we observed that in case of insertions the inserted sequence could be located at the start or end of assembled contigs, resulting in a clipped alignment to the genome. We thus also included the option to include such clipped alignments to identify the position of the insertion and at least the start or end of the inserted sequence.

Figure S2 shows an evaluation of different thresholds on the indel length with and without inclusion of clipped contig alignments for insertion detection. This showed that all indels could be detected only with a relatively small minimum indel length of 16 nt and the inclusion of clipped contig alignments. Higher minimum indel lengths excluded the 16 nt insertion in the ΔICP4 mutant, while the lacZ gene insertion in the ΔICP27 mutant would be missed without allowing clipped contig alignments. However, this parameter combination resulted in large numbers of predicted insertions for ΔICP0, ΔICP22, ΔICP27, and Δ*vhs* infection, making it difficult to distinguish the correct indels in these mutants. These results show that *de novo* assembly alone is not sufficient to identify indels with high precision without further post-processing and tuning parameters to a particular sample. In contrast, one parameter combination for our pipeline recovered all variants introduced into the null mutants without predicting too many additional indels. Notably, the additional indels identified by our pipeline for the HSV-1 null mutants were not actually incorrect as they represented indels in the parental strains of the null mutants or introns.

While a larger number of additional insertions was identified for the GFP-expressing HSV-1 virus, analysis of the identified consensus sequences of insertion starts and ends determined from clipped reads allowed easily ruling these out. A further advantage of our approach is that assembly is optional if one is not interested in the insertion sequence or the insertion is short enough that the sequence can be identified directly from the consensus sequences as in the case of the ΔICP4 mutant. Without assembly, the pipeline runs in a few minutes instead of *>* 1h with assembly, reducing the runtime enormously.

## Discussion

In this article, we present a pipeline to support functional genomics analysis of viral variants by allowing the direct identification of SNPs and indels from the functional genomics experiments, such as RNA-seq, ATAC-seq or others. Development of the pipeline was motivated by the observation that commonly used null mutants are often poorly described. Notably, this does not only apply to null mutants created decades ago before the availability of viral genome sequences but also more recently created virus variants as the GFP-expressing HSV-1 virus. In the latter case, only the approximate location relative to viral genes was described. In this way, we provide the first annotation of the precise genome location for key indels in several widely used HSV-1 variants.

Our pipeline has the advantage that it does not require additional genome sequencing experiments and can be performed directly for the experiment from which biological conclusions are drawn. Furthermore, the computational overhead is relatively low, in particular if sequence assembly for identification of (longer) insertion sequences is omitted. Despite the additional overhead, identification of inserted sequences from read assemblies has the additional advantage that it allows confirming the insertion of particular marker genes like GFP or lacZ and distinguishing the key insertions from other insertions that may have been correctly or incorrectly called.

A disadvantage of our pipeline is that it depends on sufficient read coverage of the corresponding genome regions. While this also applies to standard genome sequencing, functional genomics data can have low read coverage either in parts or on the complete genome if they depict gene expression (such as RNA-seq, PRO-seq or similar methods to capture transcriptional processes) or if the viral genome shows generally low coverage. In the first case, lowly expressed genes or non-transcribed genomic regions may have low coverage, even though most parts of viral genomes are generally transcribed to some degree. The second case applies when virus genome replication and transcription are impaired, such as during ΔICP4 and TsK infections, or in the early stages of infection. Nevertheless, this issue can be addressed by combining different types or replicates of functional genomics data or different time points of infection.

Although we only tested the pipeline for (variants of) RNA-seq data, these represented both the major challenges for our pipeline, i.e. variable and low coverage samples, and the most commonly applied assay for functional studies of viral null mutants. We are thus confident that our pipeline will be highly useful for researchers using functional genomics to study viruses and the functional role of individual virus genes.

## Acknowledgments

This work was funded by the Deutsche Forschungsgemeinschaft (DFG, German Research Foundation, www.dfg.de) in the framework of the Research Unit FOR5200 DEEP-DV (443644894) project FR 2938/11-1 to C.C.F.

## Supplementary Figures

**Figure S1.**
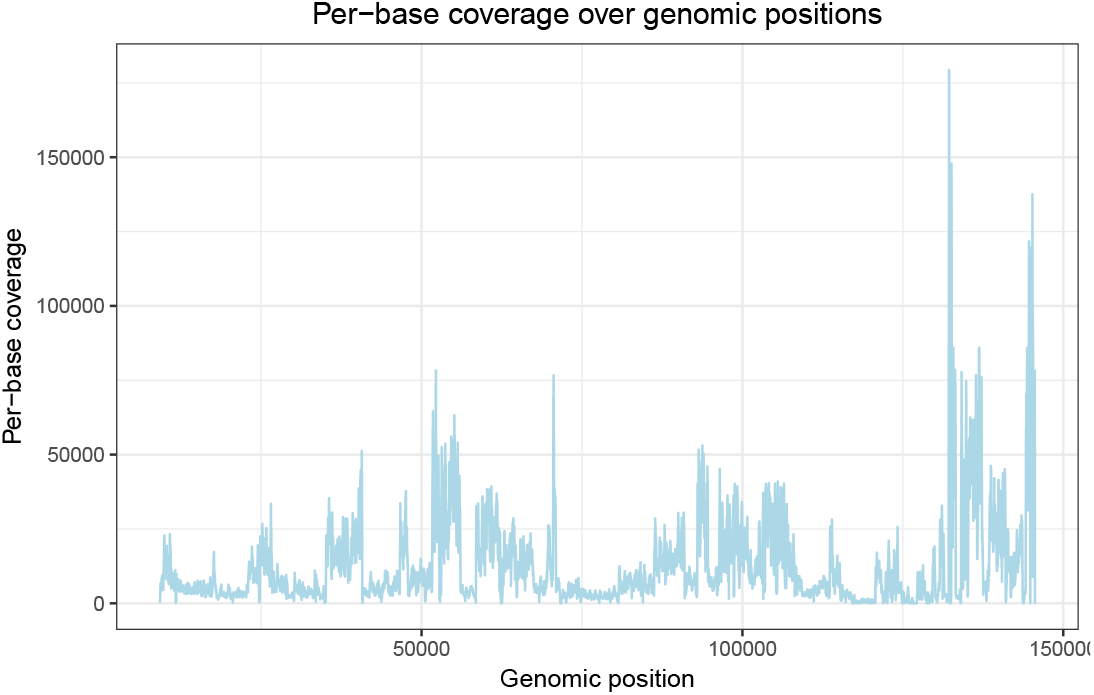
Per-base read coverage (y-axis) for HSV-1 genomic positions (x-axis) for an 4sU-seq sample for ICP22-null HSV-1 infection. This shows that read coverage varies considerably across the genome depending on gene expression.

**Figure S2.**
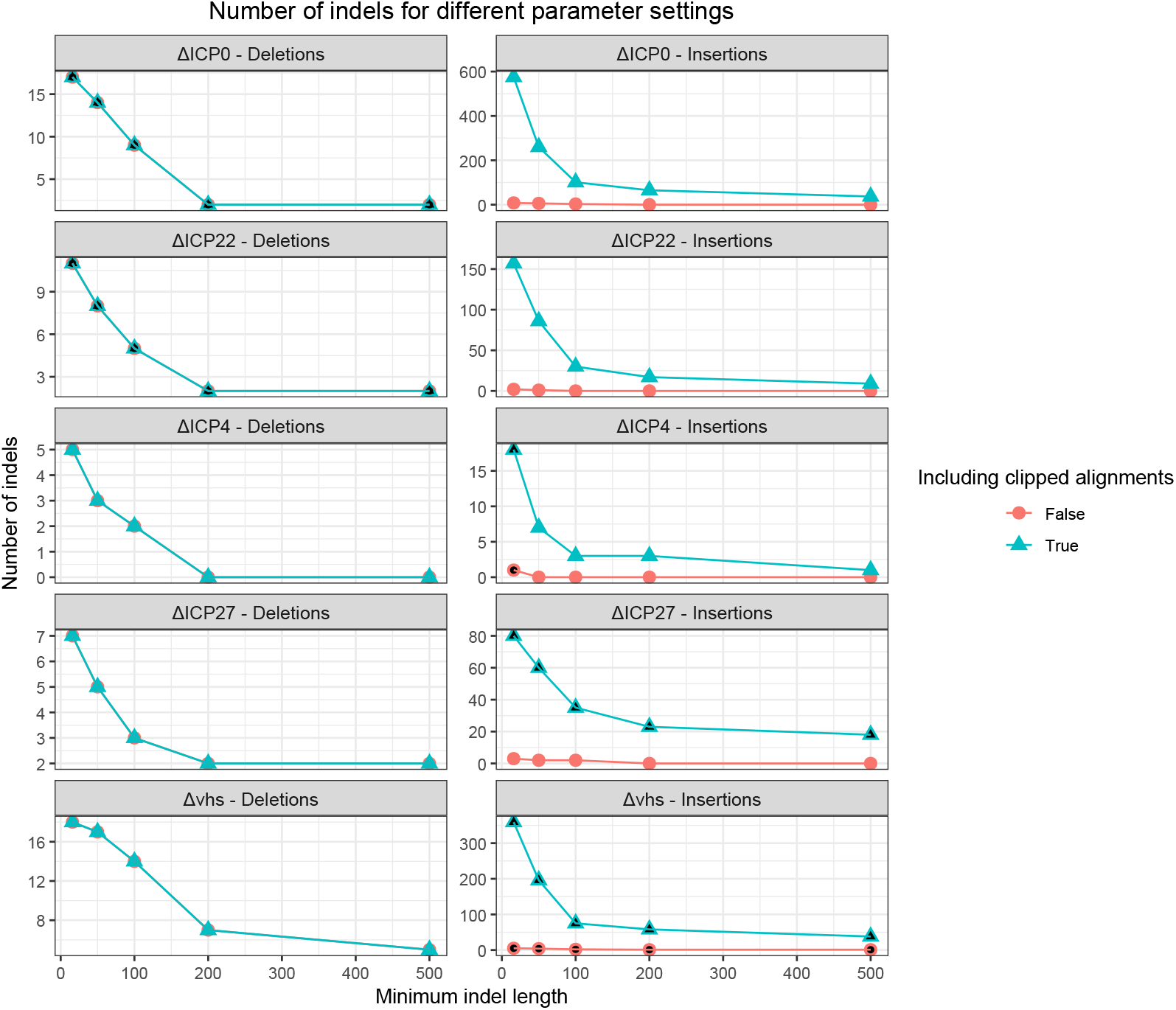
Analysis of the number of predicted deletions or insertions identified from the contigs assembled from the raw sequencing reads for different minimum indel lengths. For insertions, we also evaluated the effect of predicting insertions if only one end of the contig can be aligned to the genome in a clipped alignment. Parameters for which the correct deletion (for ΔICP0 and ΔICP22 infection) or the correct insertion (for ΔICP4, ΔICP27 and Δvhs infection) is also identified are filled in black.

